# Imputation strategy for population DNA methylation sequencing data

**DOI:** 10.1101/2025.03.21.644107

**Authors:** Alexandre Duplan, Shannon Brandt, Abel Garnier, Jorg Tost, Leopoldo Sanchez, Ludovic Duvaux, Stéphane Maury, Harold Duruflé

**Affiliations:** INRAE, ONF, BioForA, UMR 0588, 45075 Orléans, France; Université d’Orléans, INRAE, P2e, EA 1207 USC 1328, 45067 Orléans, France; INRAE, Université de Bordeaux, BIOGECO, F-33610 Cestas, France; Centre National de Recherche en Génomique Humaine, CEA-Institut de Biologie, François Jacob, Université Paris-Saclay, F-91000 Evry, France

**Keywords:** SeqCapBis, Omics, Population, epigenetics, Missing data

## Abstract

**Background:** DNA methylation is a central epigenetic mechanism involved in regulating gene expression and responses to environmental factors. Although it can sometimes be passed down through generations, its heritability remains variable depending on the species and biological context. These characteristics make it a key marker for studying genotype-environment interactions. However, whole-genome sequencing for DNA methylation analysis remains costly when applied to large numbers of individuals, prompting researchers to focus on specific regions of interest. This targeted approach often results in data matrices with missing values for some individuals, which can hinder downstream analyses.

**Results:** Our study used 200 and 189 poplar and oak individuals from natural populations, respectively. We tested and compared seven methods for missing data imputation in the specific context of targeted DNA methylation sequencing data obtained in the three different DNA methylation contexts in plants (CpG, CHG, and CHH). The comparison of the different imputation result allows to evaluate their performance to determine the most suitable approach for this type of data. Among them, NIPALS, MissForest, and LOESS provided the highest accuracy. NIPALS delivered the best overall performance but with moderate computational cost, MissForest achieved similar accuracy with faster computation, and LOESS offered competitive results suitable for large datasets.

**Conclusions:** Our results provide a reference for the selection of imputation strategies in targeted sequencing studies, improving the reliability of DNA methylation analyses and broadening the applicability of this type of data in epigenomic research.

**Graphical abstract:** 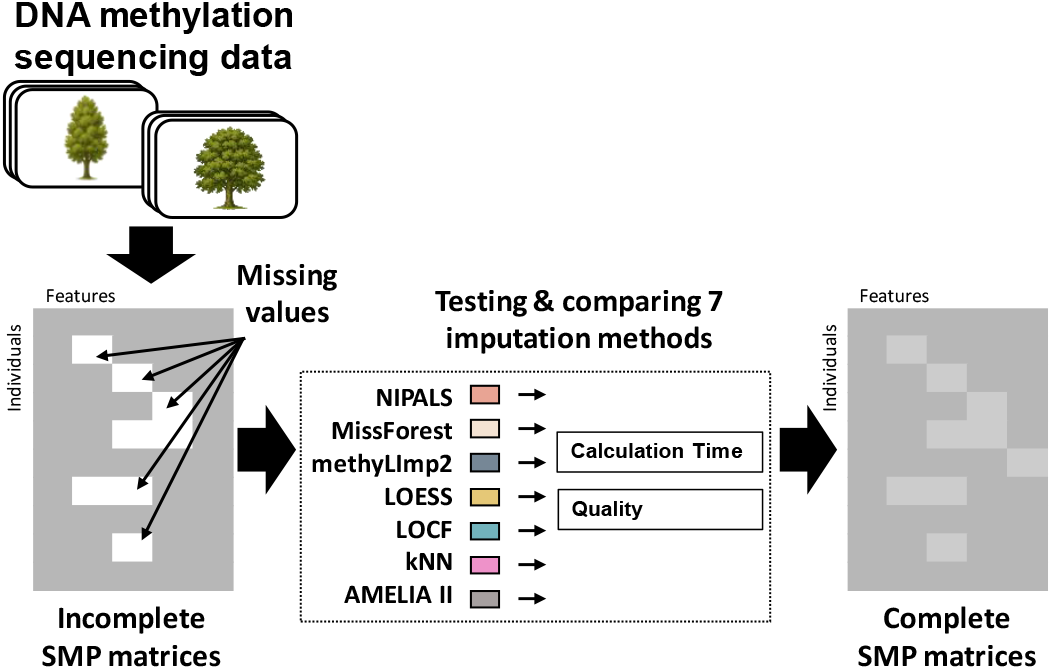

## Background

Collected datasets often contain some degree of missing or incomplete data, which can arise for various reasons. Managing missing data remains a significant challenge, and researchers and statisticians have long emphasized the importance of avoiding missing values whenever possible [1]. It is generally accepted that when a dataset has a low percentage of randomly distributed missing data, typically less than 15%, the missing values can be removed without significantly impacting the final analysis results [2, 3]. However, when the missing rate exceeds this threshold, the treatment of these missing data must be done carefully [4]. Furthermore, in many cases, even a small amount of missing data, present only in a subset of individuals, may hold valuable information that should not be overlooked. Moreover, these missing data may be distributed in a scattered manner, making the elimination decision difficult. It is acknowledged that there is no ideal method for handling missing data [5] and that understanding the reasons behind missing data is crucial before applying any imputation technique.

Large omics datasets can now be obtained with relative ease. However, their management and pre-processing are not straightforward and often requires careful consideration. Among the various omics techniques available to study biological systems, epigenomics is a more recent and stands out as particularly valuable one [6, 7]. Epigenomics focuses on stable yet reversible chemical modifications to nucleotide bases or histones, which influence gene expression without altering the underlying DNA sequence and that can be inherited. This pseudo-stability makes epigenetic modifications a key factor in understanding causal biological mechanisms under environmental conditions [8]. Epigenetics encompasses three main types of modifications: DNA methylation, histone modifications and variants and non-coding RNAs. DNA methylation, specifically, involves the reversible addition of a methyl group to the 5’ position of cytosines, a process realized by DNA methyltransferase and demethylase enzymes. In plants, DNA methylation takes place in three different contexts [7]: CpG, CHG, and CHH (with H representing the bases A, T, or C).

For DNA methylation analysis (see review of Agius *et al*., 2023), whole genome bisulfite sequencing (WGBS) is a powerful method to interrogate methylomes at single-base resolution [10, 11]. However, such comprehensive measurements are expensive for medium and large genomes, making it challenging for population-scale studies [12]. To overcome this, most published methylomes are sequenced below the saturation threshold, leading to a large proportion of cytosines with missing data or insufficient coverage. Specific methods have been developed to address this type of data. For example, METHimpute [13] is a hidden Markov model-based algorithm that enables the construction of complete methylomes by estimating the methylation state and level of all cytosines across the genome, regardless of coverage depth. BSImp [14] is a probabilistic-based algorithm that utilizes information from neighbouring sites to quickly reconstruct partially observed methylation patterns. BoostMe [15] is also an imputation method that uses a gradient boosting algorithm named XGBoost, and exploits information from multiple samples of WGBS data for prediction. All these approaches aim to accurately infer methylomes from low-coverage data (*e*.*g*., 5X), to achieve a level of accuracy comparable to high-coverage data (*e*.*g*., 50X). DNA methylation BeadChip data is another widely used type of epigenetic data, especially popular for profiling large-scale and population-based studies. For example, the Infinium HumanMethylationEPICv2 was developed for the purpose of studying the human DNA methylome [16]. Methods for imputing missing values from methylation matrices were thus developed based on linear regression, such as methyLImp [17] and methyLImp2 [18]. However, other data acquisition methods exist for DNA methylation. Targeted capture bisulfite oligonucleotide sequencing (SeqCapBis) context uses hybridization probes to capture and enrich relevant regions of interest [19]. This approach has been proposed to study the variation of DNA methylation in natural tree populations [12]. First, highly variable DNA methylation regions were identified using WGBS on a subset of genotypes that represent the genetic diversity of the population. Then, variations in DNA methylation within these targeted regions were captured and sequenced at the population level.

Currently, advanced imputation approaches can handle heterogeneous variables, but this often requires substantial computational resources. Additionally, not all imputation methods are appropriate for all data types, especially when dealing with DNA methylation data. However, a number of pan-epigenome-scale methods, such as epigenome-wide association study, require large sample sizes and no missing data. The challenge is therefore to be able to impute a high proportion of single methylation polymorphisms (SMPs) with few missing values in order to limit the loss of information. In this context, we evaluate several imputation methods on targeted DNA methylation sequencing data for two species, considering the trade-off between accuracy and computational efficiency. The comparative analysis is applied to hundreds of genotypes, from natural populations, of two different tree species, poplar and oak, using the three methylation contexts (CpG, CHG and CHH).

## Methods

### Plant material

A set of 200 *Populus nigra* genotypes from various natural populations, representative of the natural west European range of this species, were used. The trees were cultivated for four years in a common garden in Orléans (France, 47°50’06.0”N 1°54’39.6”E), as described in previous studies [20–22]. Young differentiating xylem and cambium tissues were harvested from approximately 80 cm-long stem sections, taken 20 cm above ground level and extending up to 1 m. Samples were collected from two replicates of each genotype in two blocks of the common garden design (described in Chateigner *et al*., 2020). To ensure sufficient material quantity, the DNA from both replicates was pooled before being sequenced.

For oak, the dataset consisted of 189 *Quercus petraea* genotypes from 22 populations, spanning a South-west to North-east gradient across France and Germany. These trees were cultivated in a common garden experiment located in Sillégny (France, 48°59’24.0”N 6°07’55.2”E) as described in Ducousso *et al*., 2022. Bud samples from each tree were harvested at the end of ecodormancy [12].

### Genomic DNA extraction and targeted methylated sequence capture

The methods for genomic DNA extraction have been described in the previous studies [24] and the methods for sequencing analyses have been previously described in detail [12] with a step-by-step bioinformatics manual available in the public repository protocols.io at dx.doi.org/10.17504/protocols.io.8epv5xw4ng1b/v1.

Briefly, a subset of individuals (20 for poplar and 10 for oak), considered representative of the entire population, was selected and subjected to whole-genome sequencing (WGS), enabling the detection of SNPs. Subsequently, WGBS was performed to identify regions with high methylation variation across populations. Finally, all individuals in the population were sequenced using targeted DNA methylation sequencing (SeqCapBis) approaches to focus on methylation-variant regions that were not caused by SNPs. Each methylation context was considered separately, and methylation was quantified in 1 kb sliding windows across the entire reference genome. Each window and its associated methylation level were considered as potential candidate regions for targeting, with only regions showing high levels of differential methylation between populations being used for further analysis, as previously described in detail [12]. A set of 120 bp probes was selected to capture 18 Mb of the genome. All the capture experiments were performed at the PGTB and sequencing was performed at the GeT-PlaGe facility on an Illumina NovaSeq 6000. Mapping was performed with BSMAPz 1.1.3 [25] against the *Populus trichocarpa* V4.1 reference genome for poplar [26] and the *Q. robur* reference genome (Haplome V2.3) for oak [27]. DNA methylation analysis was performed using the methylKit package 1.18.0, and the data were filtered to retain only sites with at least 5X coverage [28].

### Benchmark datasets and missing data

To limit calculation times for imputation analysis, only chromosome 16 of the poplar genome was used because of its average size. This chromosome has a size of 14,802,566 bp [26] including in this study 665,168 of single methylation polymorphisms (SMP) distributed in 59,070 CpG (including 12,957 without missing data), 97,748 CHG (including 27,023 without missing data), and 508,350 CHH (including 120,025 without missing data) along the chromosome.

For oak, calculation time was also limited by using a subset of the oak genome. The CpG SMP dataset was split into 20 similarly sized bins, ensuring that known scaffolds were not split. In this study, Bin 1 is used as it contains the least amount of scaffolds (Sc0000001, Sc0000047, Sc0000011, Sc0000017, Sc0000003, Sc0000027, Sc0000005). This subset contains 443,687 SMP distributed in 42,504 CpG (including 6,509 without missing data), 55,272 CHG (including 7,230 without missing data) and 345,911 CHH (including 33,024 without missing data).

The value of an SMP is defined as the ratio of the number of reads showing a methylated cytosine to the total number of reads at a specific cytosine position in the genome. This value corresponds to a β-value, therefore ranges from 0 (unmethylated) to 1 (methylated), reflecting the degree of methylation at that particular position [29].

The poplar dataset has fewer missing data than the oak dataset (Fig. 1). In detail, for poplar, 12,957 CpG SMPs (22%), 27,023 CHG SMPs (28%), 120,025 CHH SMPs (24%) had no missing data across all 200 genotypes. We observed 22,304 CpGs (38%), 43,747 CHG (45%), 204,471 CHH (40%) with 10 or less missing values across all 200 genotypes. For oak, 6,509 CpG SMPs (15%), 7,230 CHG SMPs (13%), 33,024 CHH SMPs (10%) had no missing data across all 189 genotypes. We observed 12,412 CpGs (29 %), 15,197 CHG (27 %), 73,974 CHH (21%) with 10 or less missing values.

**Figure 1.**
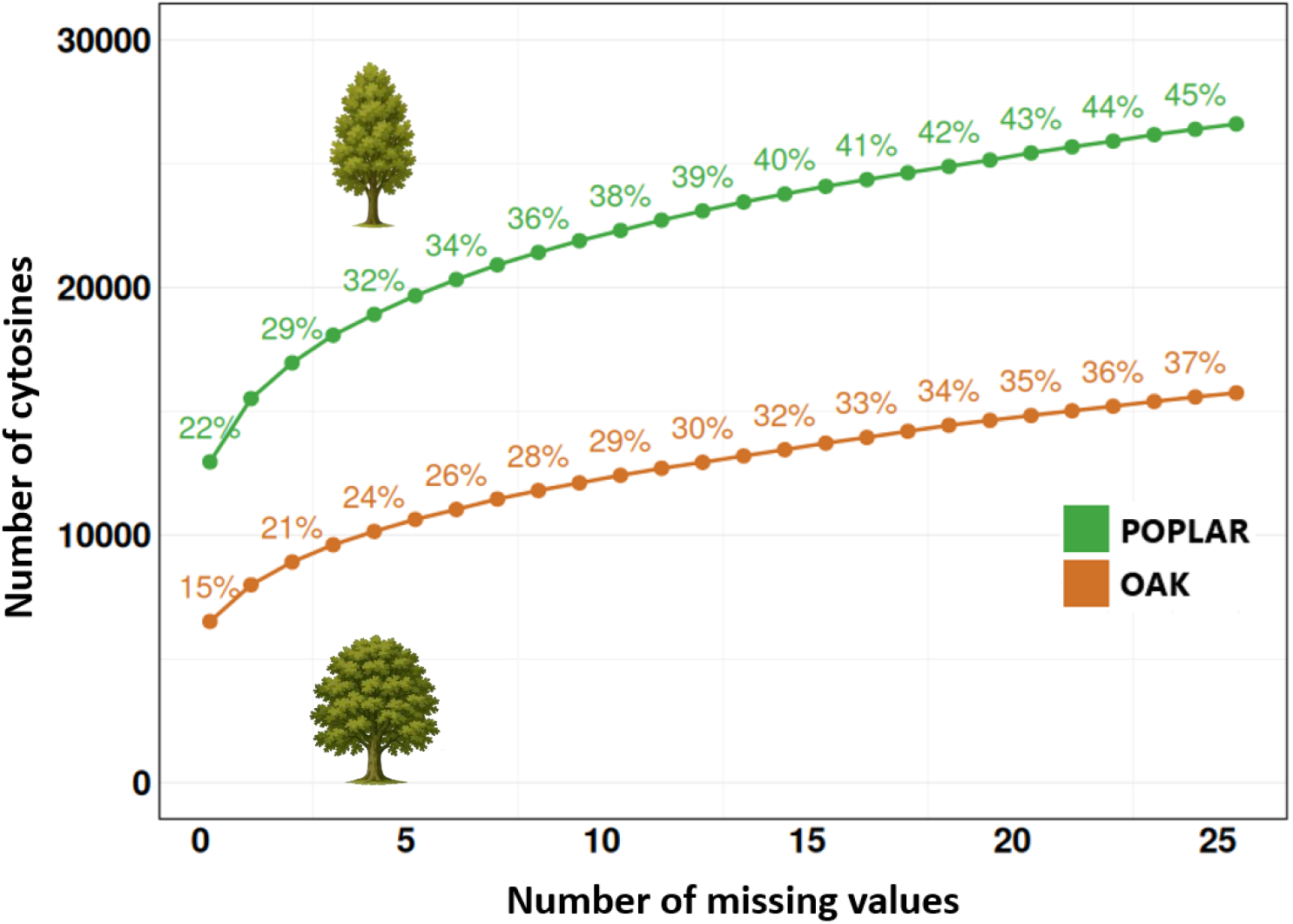
Cumulative number of SMPs per missing values in CpG context of the chromosome 16 of poplar and the Bin 1 of oak.The percentage shown above the curves indicates the proportion of SMPs captured across all data within the CpG context.

Methylation ratios in the CpG context exhibit high variability, with values ranging from 0 to 1 (Fig. S1) for both species. In the CHG context, variability is comparable to that of CpG, with a greater number of SMPs in this dataset. The CHH context shows lower variability, with values most often ranging from 0 to 0.2, but this dataset contains much more data, with approximately ten times the number of SMPs compared to the other contexts.

The density distribution of SMPs with respect to the number of missing values reveals a distinct pattern across both species and methylation contexts (Fig. S2 & S3). About a quarter of SMPs exhibit low levels of missing data (*i*.*e*., concerning less than 10 individuals). Finally, another peak in the distribution highlights a subset of SMPs with a high number of missing values. For example, in the case of the oak, 5,853 SMPs were found in CpG, 7,427 in CHG, and 53,686 in CHH that were detected in fewer than 10 individuals. Similarly, for the poplar, 9,075 SMPs in CpG, 13,156 in CHG, and 70,225 in CHH were identified under the same criteria.

### Computational information

All statistical and genetic analyses were run using R v4.4.0 [30] and analyses were carried out on the Genotoul Bioinformatics platform (http://bioinfo.genotoul.fr). Concerning the resources explicitly requested in our SLURM bids. The analyses were carried out using SLURM job submissions with CPU resources set at 10 and 100GB RAM.

### Imputation Algorithm

#### Amelia II

This approach performs multiple imputations of missing observations simultaneously using an EMB Expectation-Maximization with Bootstrapping algorithm. This algorithm is designed to be faster and more flexible than Markov Chain Monte Carlo based methods [31]. Multiple imputations serve to create several independent versions of the dataset, each with different imputed values. This allows for the estimation of the variability and uncertainty introduced by the missing data, providing more robust and reliable results. The algorithm is particularly efficient in handling large and complex datasets with missing values that may not be missing at random [32]. For this approach, the R package `Amelia` v1.8.3 was used [31], with the parameter M set to 3. This parameter specifies the number of imputations to generate and sets it to 3 to maintain a realistic calculation time.

#### k-nearest neighbour (kNN)

The kNN method [33] replaces a given missing value with the mean of the nearest neighbours, based on a similarity measure of the observations of that same variable. Distance functions can thus be used to select neighbours to allow for the inclusion of numerical variables. An aggregation of the k values of the kNN is used as the imputed value. For this study, the parameter for the number of (k) neighbours was fixed at 8. The calculation of the distance to define the nearest neighbours is based on an extension of the Gower distance [34], where the distance between two observations is the weighted average of the contributions of each variable. The aggregation of the k values into an imputed value is the default median for continuous variables. The main advantages of the kNN model are: 1) imputed values are real values and not constructed; 2) it uses auxiliary information provided by the values themselves, which preserves the initial structure of the data; 3) it is non-parametric and therefore does not require the specification of a predictive model [35]. For this approach, the R package `VIM` v6.2.2 was used [36]. To reduce calculation time while providing a preliminary assessment of the model’s performance, the evaluation of the kNN imputation model in the context of CHH methylation was carried out on a subset of the first 15,000 SMPs. The parameters choice of K = 8 was based on preliminary performance evaluations, as shown in Figure S4.

#### Last Observation Carry Forward (LOCF)

When an observation is missing for a sample, it is replaced by the last available observation for that same sample. The method assumes that the result remains constant at the last observed value after the observation [37]. For this approach, the R package `zoo` v1.8.12 was used [38].

#### LOcal regrESSion (LOESS)

Local regression imputes missing data using a low-degree polynomial fitted around the missing data by weighted least squares, giving more weight to values close (*i*.*e*. a neighbourhood in the space of independent variables) to the missing data [39]. For this approach, the R package `locfit` v1.5.9.10 was used [40] as described in [41].

#### methyLImp2

methyLImp is a missing value imputation method developed for DNA methylation BeadChip data [17]. This algorithm is based on solving many multiple linear regressions with the aim of improving the observed inter-sample correlation at cytosine methylation levels. This method allows parallelizing the imputation by chromosomes in order to impute missing values using only probes on the same chromosome for the CpG context. For this approach, the R package `methyLlmp2` v1.0.0 was used [18].

#### MissForest

This approach is a random forest imputation algorithm for missing data. For each variable with missing values, the algorithm fits a random forest on the observed part using other variables as explaining variables before predicting the missing part. This training and prediction process repeats iteratively until a stopping criterion is met or a user-specified maximum number of iterations is reached. The reason for these multiple iterations is that, from iteration 2 onwards, the random forests performing the imputation will be trained on higher quality data. For this approach, the R package `missForest` v1.5 was used [42], with the parameters maxiter = 10 and ntree = 10, to limit calculation time. The parameters choice was based on preliminary performance evaluations, as shown in Figure S5.

#### Nonlinear Iterative Partial Least Squares (NIPALS)

The NIPALS algorithm [43] is an iterative Jacobi-type method that estimates the elements of a Principal Component Analysis (PCA) of a finite-dimensional random vector. This can be done even in the presence of missing data, without the need to remove the affected individuals or impute these missing values [44], making it a method specifically adapted to this type of data [45]. NIPALS then uses the components estimated by PCA to reconstruct the missing elements in the data. In this study, we set the default parameter (ncomp = 10), which specifies the number of principal components to extract from the data. For this approach, the R package `mixOmics` v6.28.0 was used [46].

#### Performance evaluation

To assess imputation quality, we used three standard metrics: Root Mean Squared Error (RMSE), Mean Absolute Error (MAE), and the coefficient of determination (R^2^). RMSE quantifies the square root of the average squared differences between the imputed and true values. It is sensitive to large errors and emphasizes substantial deviations. MAE calculates the average absolute difference between imputed and true values. It provides a more robust and interpretable estimate of average error, being less influenced by outliers than RMSE. The R^2^ coefficient measures the proportion of variance in the true values that is captured by the imputed values. An R^2^ close to 1 indicates that the imputation closely matches the original data, whereas an R^2^ near 0 suggests little to no improvement over simply predicting the mean.

## Results

In this study, we compared the performance in terms of accuracy and calculation time for seven single methylation polymorphism (SMP) imputation methods using targeted DNA methylation sequencing data. The benchmark was carried out by using data from the three methylation contexts present in plants (CpG, CHG and CHH) for two sets of natural populations of poplar and oak.

We evaluated the performance of the seven imputation methods described above using the benchmark datasets in poplar and oak. To do this, we started with the largest subset of data with no missing values in each species, which represented 22% and 15% of the total SMP in poplar and oak, respectively (Fig. 1). On this basis, to create our test datasets, we randomly selected and masked 30% of the positions (β-values) of 10 different individuals, with individuals re-sampled independently at each position. This process was performed 12 times to generate 12 different alternative datasets for each species. Imputation performances were calculated as the correlation between the imputed data and the actual data in each dataset and for each method (Fig. 2).

**Figure 2.**
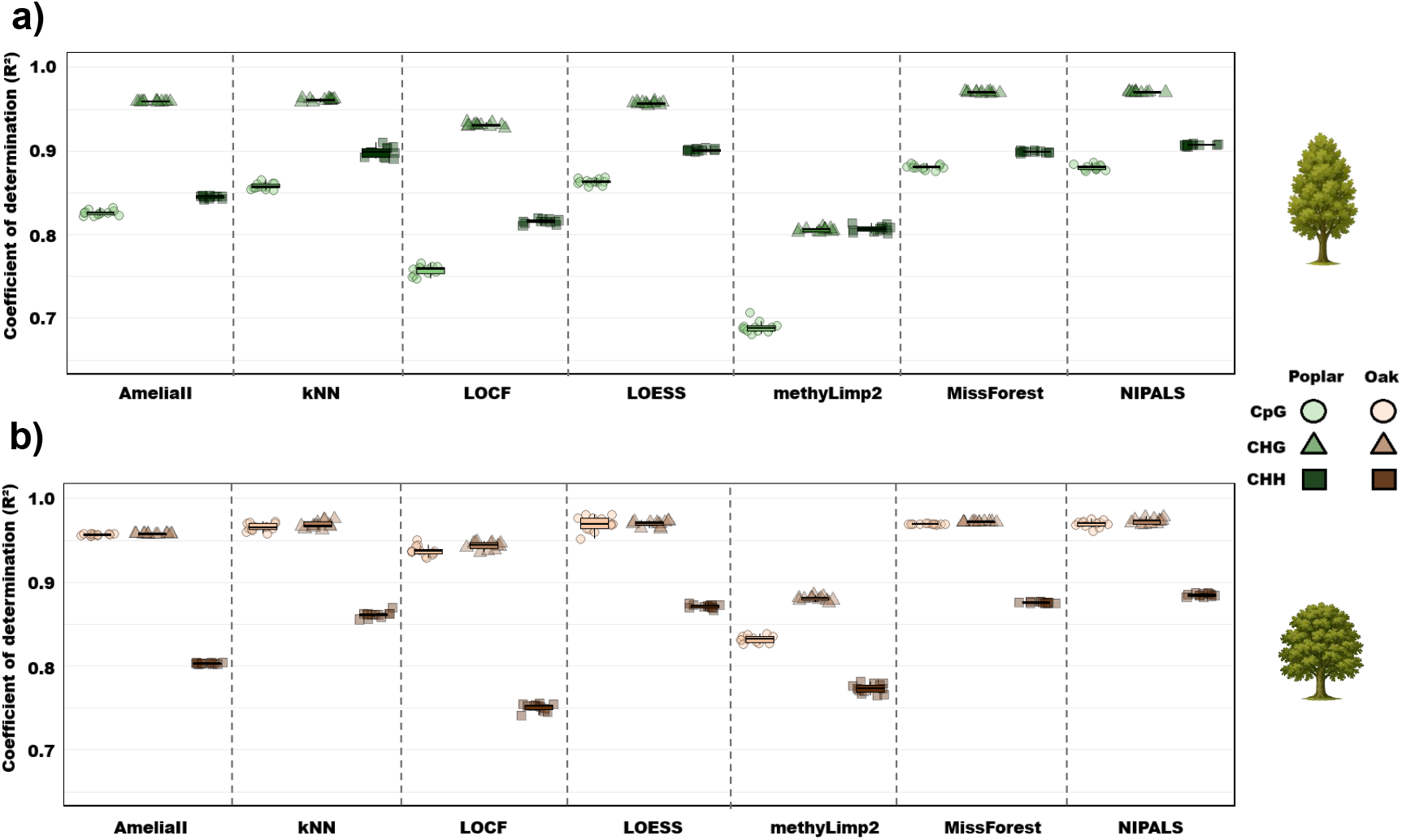
Comparison of the results of seven imputation models using the coefficient of determination (R^2^) between imputed and real data for a) Chromosome 16 in poplar b) Bin 1 in oak. In both cases, 30% of SMPs have 10 missing data points across 12 independent tests.

Overall, the imputation result showed a trend towards higher coefficient of determination (R^2^) in the CHG methylation context compared to the other contexts, with an average of 0.936 and 0.952 for poplar and oak, respectively. The CpG datasets show lower R^2^ than CHH for poplar and the opposite for oak, where the R^2^ for CpG is just as high as CHG. The range of R^2^ values varied between 0.75 to 0.97 for all methods and data types except for the methyLImp2 method, which showed R^2^ values of 0.68 for poplar in CpG context. R^2^ values were all relatively high across the various methods, with little variation between the 12 replicates.

The NIPALS imputation method showed the highest R^2^ values across all three contexts, for both species datasets, in poplar (CpG: 0.880, CHG: 0.970, CHH: 0.907) and in oak (CpG: 0.970, CHG: 0.973, CHH: 0.885). The methyLImp2 algorithm had the highest MAE and RMSE value and the lowest R^2^ value in three contexts for both species (Table 1) except for oak in the CHH context, where it ranked second lowest (0.773) with the LOCF method showing the lowest R^2^ (0.751). For the poplar datasets, NIPALS and MissForest showed the highest R^2^ of 0.880. Moreover, MissForest shows the lowest RMSE value (0.146) followed by NIPALS (0.147). Concerning the MAEs, the kNN algorithm showed the lowest value (0.081), followed by NIPALS and then MissForest (0.085 and 0.086 respectively). For oak datasets, results were similar. NIPALS and LOESS showed the highest R^2^ of 0.970, followed by MissForest (0.969). NIPALS had the lowest RMSE (0.076) followed by MissForest (0.078). Concerning the MAEs for oak, the kNN algorithm also shows the lowest value (0.041), followed by NIPALS (0.043) and MissForest and LOESS (0.045 for both). Globally, the same patterns observed in the CpG context were also observed for the CHG and CHH context in both species (Table S1 and S2).

**Table 1.** Overall CpG imputation performance was evaluated across 12 tests conducted on 200 genotypes of poplar and oak, with 30% of SMPs containing 10 missing data points.

|  |  | Calculation | RMSE | MAE | R² |
| --- | --- | --- | --- | --- | --- |
|  | Method | Time<br>(h:min:s) |  |  |  |
| Poplar | LOCF | 00:00:01 | 0.209 | 0.110 | 0.757 |
|  | LOESS | 00:00:11 | 0.157 | 0.091 | 0.863 |
|  | Amelia II | 00:02:14 | 0.177 | 0.123 | 0.826 |
|  | MissForest | 00:43:43 | 0.147 | 0.086 | 0.880 |
|  | methyLImp2 | 02:28:56 | 0.237 | 0.124 | 0.689 |
|  | NIPALS | 02:33:31 | 0.146 | 0.085 | 0.880 |
|  | kNN | 19:43:41 | 0.160 | 0.081 | 0.858 |
| Oak | LOCF | 00:00:01 | 0.113 | 0.057 | 0.938 |
|  | LOESS | 00:00:04 | 0.080 | 0.045 | 0.970 |
|  | Amelia II | 00:01:04 | 0.093 | 0.063 | 0.957 |
|  | MissForest | 00:06:22 | 0.078 | 0.045 | 0.969 |
|  | methyLImp2 | 00:10:44 | 0.183 | 0.077 | 0.832 |
|  | NIPALS | 00:46:53 | 0.076 | 0.043 | 0.970 |
|  | kNN | 02:26:04 | 0.084 | 0.041 | 0.966 |

Note: The results have been averaged over the 12 random replicas per percentage of missing data. Best results per metric are highlighted in bold for each dataset. RMSE: Root Mean Square Error. MAE: Mean Absolute Error. R<sup>2</sup> Coefficient of determination.

Concerning calculation time, the imputation models followed the order: LOCF, LOESS, Amelia II, methyLImp2, MissForest, NIPALS, and kNN, for both species, in the three methylation contexts (Table 1). In the CHH context, the times were relatively low for kNN because the dataset was reduced as too resource- and time-consuming (Table S1).

## Discussion

Because population-level DNA methylation sequencing data are specific and require specialized approaches, only a few imputation methods have been specifically adapted to these data. In this study, we evaluate seven different imputation methods to address the challenge of imputing targeted DNA methylation sequencing data for populations, for which no established operational method is currently recommended. These were evaluated and compared using the three distinct contexts of plant methylation. The targeted DNA methylation sequencing data come from two tree species, poplar and oak, and are classified according to the three methylation contexts (CpG, CHG, CHH). This distribution of missing data show that imputing SMP values for positions captured in the large majority of individuals (∼90%) can be a relevant approach to maximize the use of available data. Moreover, this suggests that some positions are widely represented in the dataset and indicates that another part appears to be specific to certain individuals or populations. Furthermore, the three methylation contexts show expected variability, with high variability in the CpG, and to a lesser extent in the CHG context, and low variability for the CHH data (Fig. S1). The amount of data is also expected, with about ten times more SMPs in CHH than in the other two contexts. To compare imputation methods, we rely on the accuracy and computation time of seven different algorithms performed on 12 independent tests, imputing 10 missing data in 30% of SMPs.

Maybe not surprisingly, LOESS algorithms are able to impute CpG DNA methylation data with high R^2^ in both species. It is well-established that neighbouring CpG sites often exhibit similar methylation levels, a phenomenon referred to as co-methylation [47–49]. However, the kNN algorithm is by far the slowest method tested here. This may be related to the calculation time needed to calculate the nearest neighbour average, making the method inappropriate for large datasets.

On the contrary, using local regressions to impute missing data in methods like LOESS or LOCF is very fast. However, the low R^2^ obtained by the LOCF method (second lowest R^2^) demonstrates that this algorithm is not suitable to impute missing data for targeted DNA methylation sequencing experiments. This limitation can be explained by the fragmented nature of the data due to the capture design. Since the sequencing does not provide continuous coverage, the last observed cytosine is not necessarily adjacent to the missing site on the chromosome. As a result, LOCF assumes spatial continuity where there is none, likely leading to inaccurate imputations and contributing to the low R^2^ observed. This strategy could be relevant for SMPs, if continuous data is used.

The Amelia II imputation method shows diverse coefficient of determination between methylation context and tree type, but it is generally the fourth and sixth most efficient model between the seven tested here. Thus, the higher RMSE and MAE scores observed at these lower R^2^, compared to other algorithms, do not encourage the use of this type of imputation method for methylation data. The random forest-based imputation method, MissForest, demonstrates superior performance in imputing SMP data, achieving high R^2^ and the low RMSE and MAE score for the three methylation contexts and in both species. However, it is the second-slowest method for poplar and requires a substantial amount of calculation time. Surprisingly, the results of methyLImp2 are largely inferior compared to the other approaches. Additionally, this algorithm displays a unique behaviour for poplar data, which contrasts with the patterns observed in the analysis using the other six methods. The methyLImp2 algorithm has been developed for missing value imputation specific for DNA methylation BeadChips for human data. We think that these results are due to the fact that this algorithm needs to be properly calibrated for targeted DNA methylation sequencing data for plants. The NIPALS imputation method showed the highest R^2^ and the lowest RMSE values in all three contexts, for both species datasets. The performance of this method is similar to kNN, while showing reasonable running times.

## Conclusion

In this study, we tested and compared seven methods for missing data imputation in the specific context of targeted DNA methylation sequencing data in plants. These methods are based on different imputation strategies, including k-nearest neighbours (kNN) averaging, bootstrapping (Amelia II), low-degree polynomial fitting (LOESS), random forest approaches (MissForest), and iterative Jacobi-type algorithms (NIPALS). Additionally, methyLImp2, which uses a linear regression approach to leverage inter-sample correlations inherent to methylation data, demonstrated strong performance on human DNA methylation matrices but was found to be unsuitable for DNA methylation targeted sequencing data in plants.

This study highlights that few methods, NIPALS, MissForest, and LOESS, achieve high imputation accuracy for methylation data, but exhibit significant differences in computational time. NIPALS offers the advantage of displaying the best imputation values while preserving the variance structure of the data. However, this method has a moderate computational cost. MissForest achieves comparable accuracy with much lower computational time, making it an attractive alternative when computational resources or time are more limited. Finally, LOESS provides competitive results and appears particularly suitable for large-scale datasets.

Given these considerations, we suggest that the choice of imputation method should be guided by the specific constraints and objectives of the study. If computational time is not a limiting factor and preserving the multivariate structure of the data is essential, NIPALS remains a sound choice. If a balance between accuracy and speed is required, MissForest offers an excellent compromise. For exploratory analyses or situations where computational efficiency is paramount, LOESS offers a remarkably fast and sufficiently accurate alternative.

Overall, this study provides a framework to guide researchers in choosing the most appropriate imputation strategy for plant DNA methylation data. However, caution is still required when interpreting biological analyses incorporating imputed values. Despite the robustness of imputation techniques, erroneously imputed values are unavoidable and can introduce bias, potentially affecting downstream analyses and biological conclusions.

## List of abbreviations

bp: base pairs
CpG: methylation context 5’-C-phosphate-G-3’
CHG: methylation context 5’-C-H-G-3’, where H correspond to A, T or C
CHH: methylation context 5’-C-H-H-3’, where H correspond to A, T or C
DNA: Deoxyribonucleic Acid
kNN: k-nearest neighbour
LOCF: Last Observation Carry Forward
LOESS: LOcal regression
MAE: Mean Absolute Error
NIPALS: Nonlinear Iterative Partial Least Squares
PCA: Principal Component Analysis
R^2^: coefficient of determination
RMSE: Root Mean Squared Error
SMPs: Single Methylation Polymorphisms
WGBS: Whole Genome Bisulfite Sequencing

## Declarations

### Ethics approval and consent to participate

Not applicable.

### Consent for publication

Not applicable.

### Availability of data and materials

The data and R Markdown are freely available online at: https://doi.org/10.57745/SUOZVE

### Author Contributions

Conceptualization: AD, HD, SM, LS.

Data acquisition: JT, AG.

Data curation: AD, SB.

Formal analysis: AD, SB.

Funding acquisition: SM.

Investigation: AD, HD.

Writing – original draft: HD, AD.

Writing – review & editing: All authors.

## Acknowledgment

Capture experiments were performed at the PGTB facility (https://doi.org/10.15454/1.5572396583599417E12) and sequencing was performed at the GeT-PlaGe facility with a special thanks to Christophe Boury and Erwan Guichoux (GeT, https://doi.org/10.15454/1.5572370921303193E12). We are grateful to the genotoul bioinformatics platform Toulouse Occitanie (Bioinfo Genotoul, https://doi.org/10.15454/1.5572369328961167E12) for providing help and/or computing and/or storage resources. We also thank GBFOR (INRAE, 2018. Forest Genetics and Biomass Facility, https://doi.org/10.15454/1.5572308287502317E12) for support for the cultivation and quality production of poplar trees. We also thank Odile Rogier and Isabelle Lesur for the development of the bioinformatics pipeline as well as Romane Callarec for the preliminary analyses carried out on the oak data. The authors are also very grateful to Mathieu Emily for his kind proofreading.

## Additional Data

**Additional file: Figure S1**. Variability of DNA methylation data on chromosome 16 in poplar and the Bin 1 in oak across the CpG, CHG, and CHH contexts.

**Additional file: Figure S2**. Density distribution of missing values per SMPs site in the CpG, CHG, and CHH methylation contexts in Bin 1 for oak and chromosome 16 for poplar. The red dotted line indicates the threshold of 10 missing values per site.

**Additional file: Figure S3**. Cumulative distribution of missing values per SMP in the CpG, CHG, and CHH contexts in Bin 1 for oak and chromosome 16 for poplar. Each curve represents the empirical cumulative distribution of the number of missing values per site for a given species and context. The red dotted line indicates the threshold of 10 missing values per site.

**Additional file: Figure S4**. Performance of the kNN imputation model across 20 different k neighbour values in CpG contexts. a) Boxplot of the coefficient of determination (R^2^ values), b) Root mean square error (RMSE), c) Mean absolute error (MAE). The evaluation was conducted on Chromosome 16 for Poplar and BIN 1 for Oak, with 12 replicates per k value and species.

**Additional file: Figure S5**. Effect of different missForest parameter setups (maxiter - number of iterations; ntree-number of trees) on CpG context of BIN1 of oak with (a) the coefficient of determination (R^2^), b) RMSE value, c) MAE value and d) the average of calculation time.

**Additional file: Table S1**. Overall CHG imputation performance was evaluated across 12 tests conducted on 200 genotypes of poplar and oak, with 30% of SMPs containing 10 missing data points.

**Additional file: Table S2**. Overall CHH imputation performance was evaluated across 12 tests conducted on 200 genotypes of poplar and oak, with 30% of SMPs containing 10 missing data points.

