## Additional files for "Imputation strategy for population DNA methylation sequencing data"

**Additional file: Figure S1.** Variability of DNA methylation data on chromosome 16 in poplar and the Bin 1 in oak across the CpG, CHG, and CHH contexts.

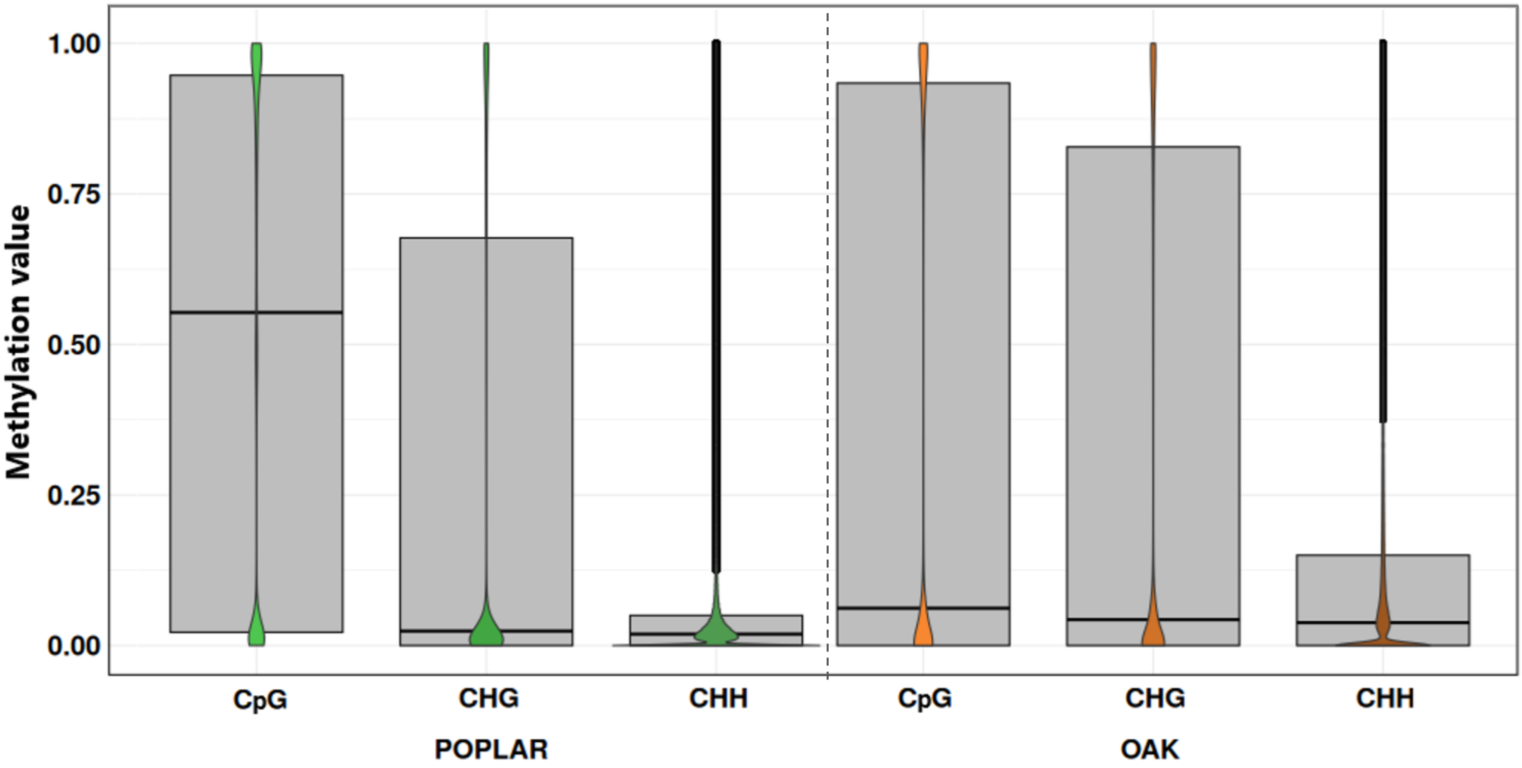

**Additional file: Figure S2.** Density distribution of missing values per SMPs site in the CpG, CHG, and CHH methylation contexts in Bin 1 for oak and chromosome 16 for poplar. The red dotted line indicates the threshold of 10 missing values per site.

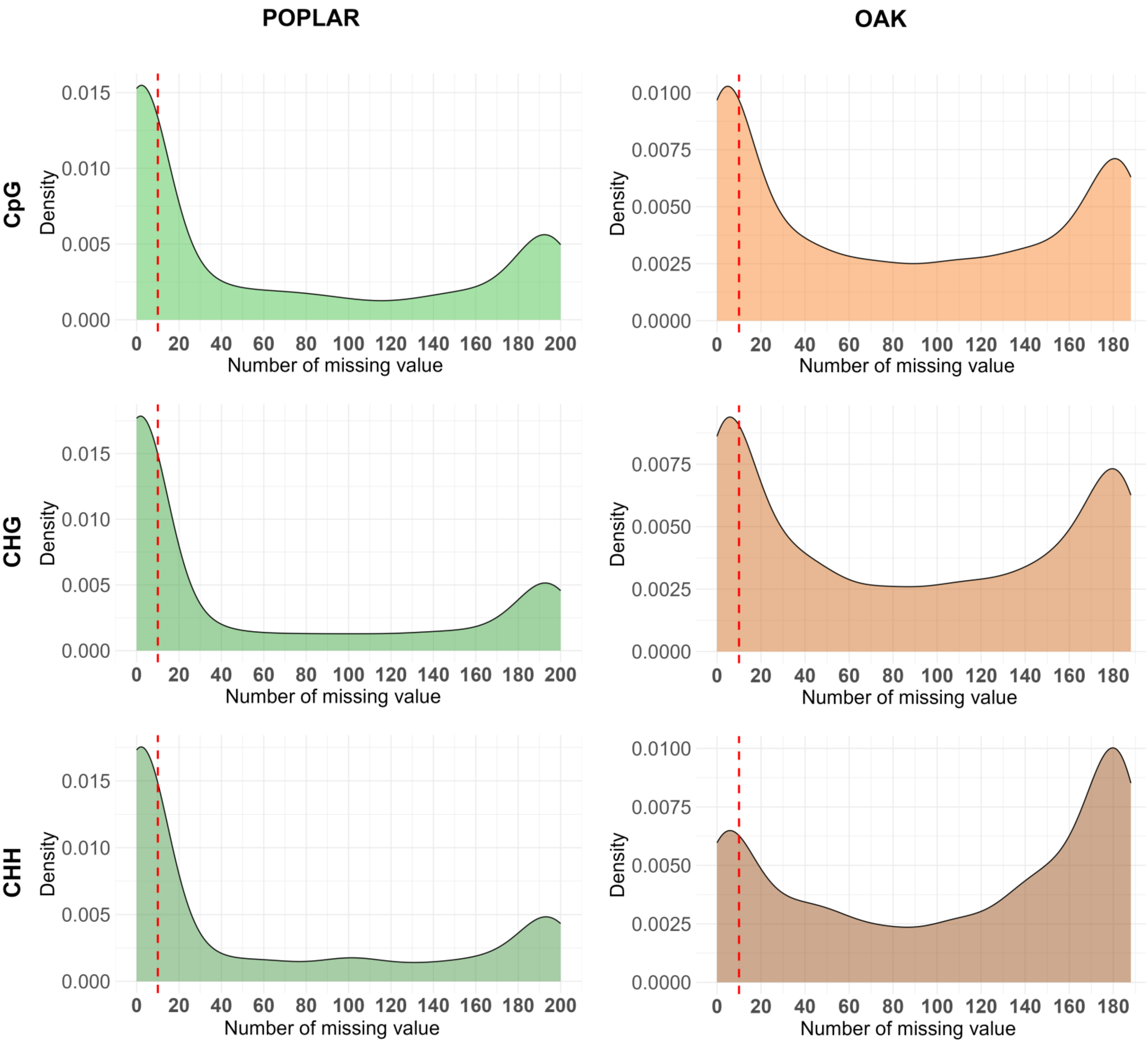

**Additional file: Figure S3.** Cumulative distribution of missing values per SMP in the CpG, CHG, and CHH contexts in Bin 1 for oak and chromosome 16 for poplar. Each curve represents the empirical cumulative distribution of the number of missing values per site for a given species and context. The red dotted line indicates the threshold of 10 missing values per site.

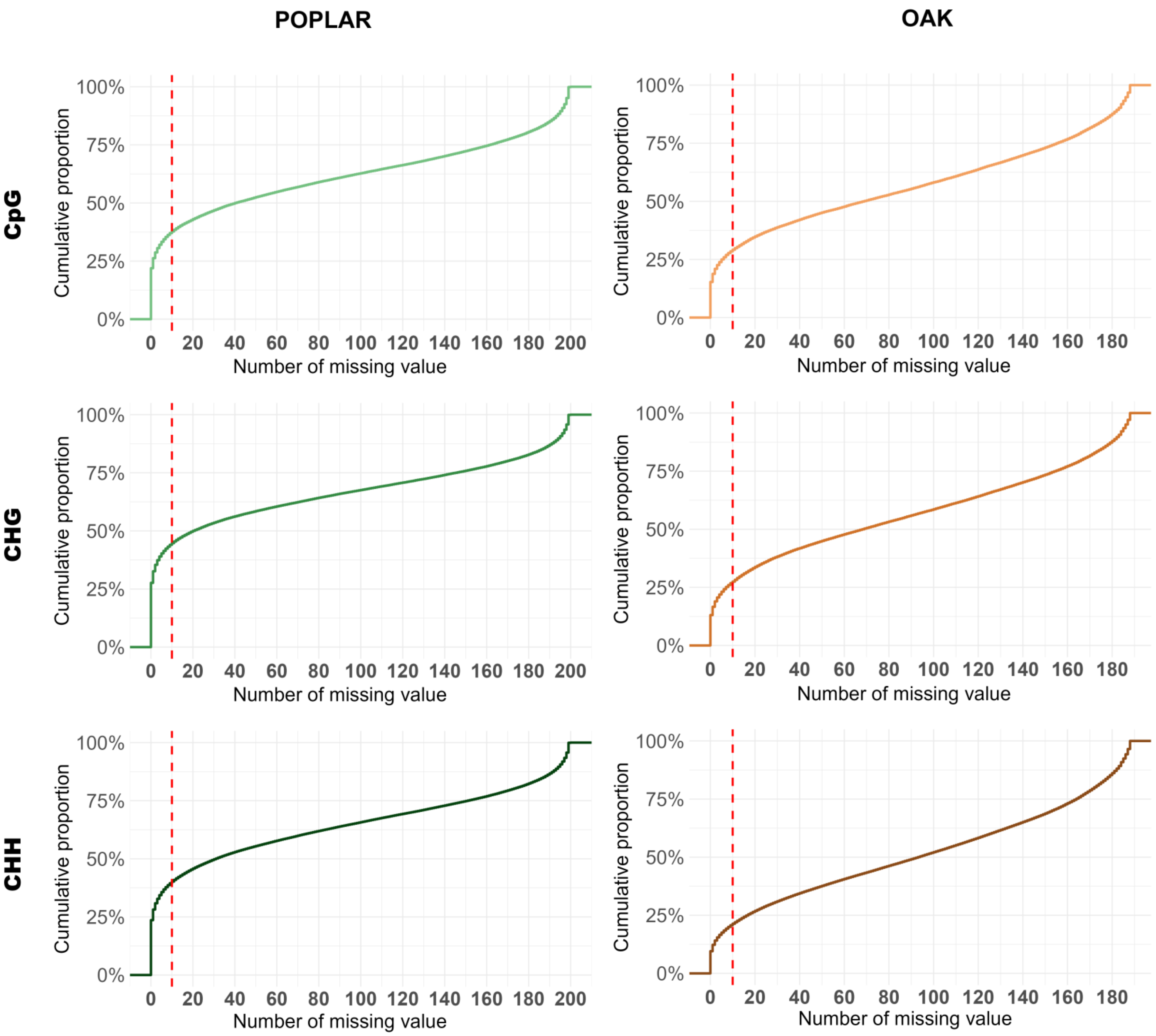

**Additional file: Figure S4.** Performance of the kNN imputation model across 20 different k neighbour values in CpG contexts. a) Boxplot of the coefficient of determination ($R^2$ values), b) Root mean square error (RMSE), c) Mean absolute error (MAE). The evaluation was conducted on Chromosome 16 for Poplar and BIN 1 for Oak, with 12 replicates per k value and species.

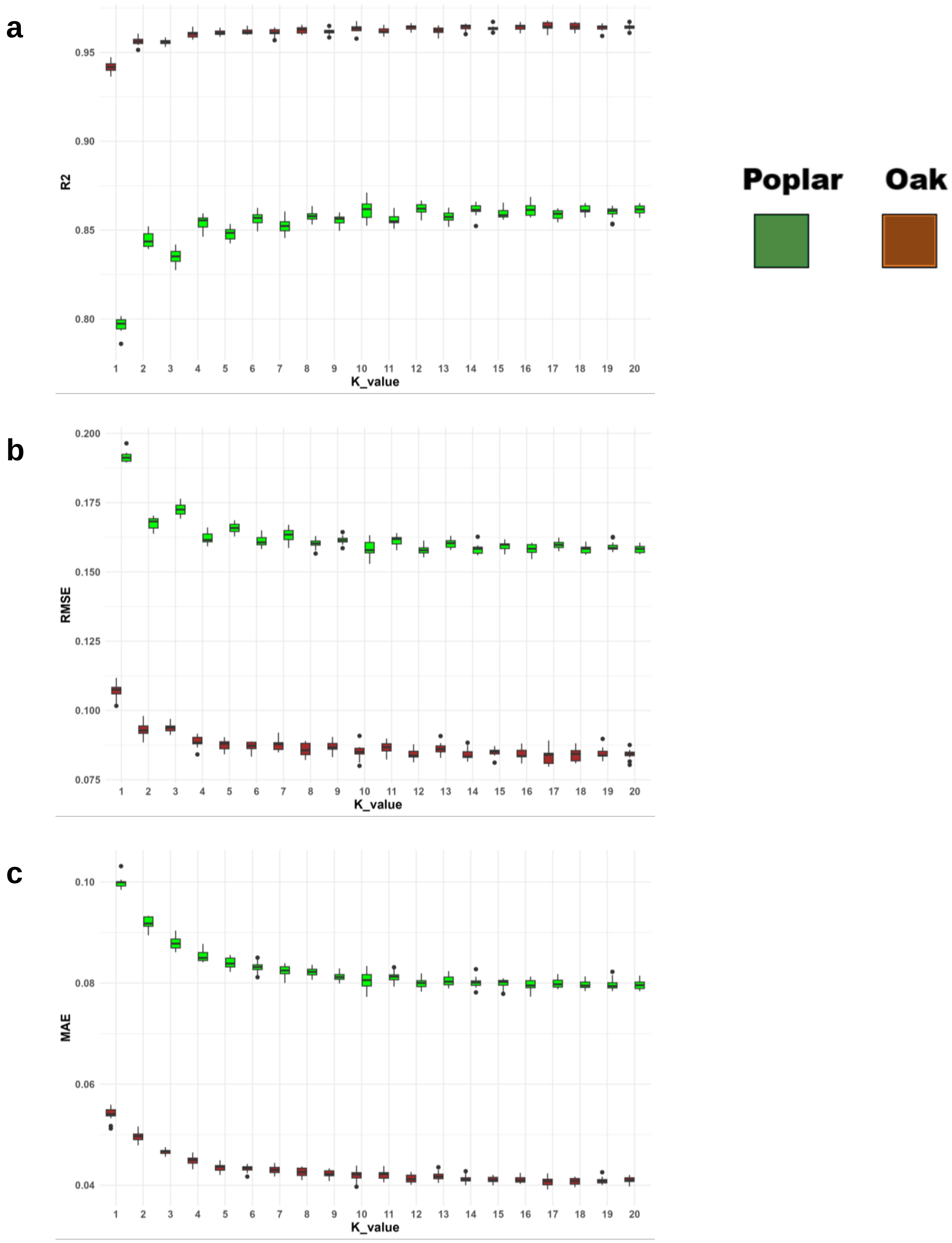

**Additional file: Figure S5.** Effect of different missForest parameter setups (maxiter -number of iterations; ntree -number of trees) on CpG context of BIN1 of oak with (a) the coefficient of determination ($R^2$), b) RMSE value, c) MAE value and d) the average of calculation time.

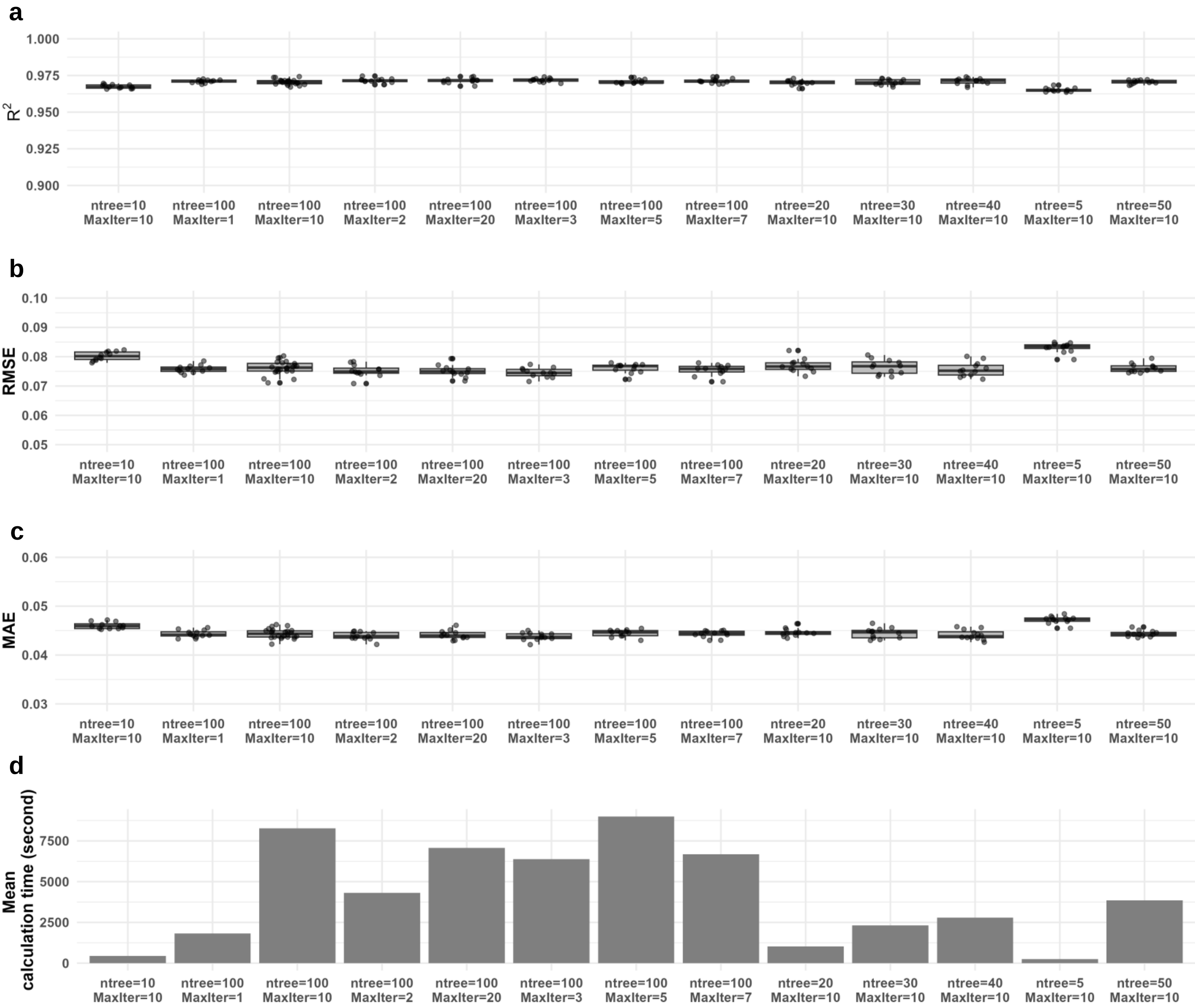

**Additional file: Table S1.** Overall CHG imputation performance was evaluated across 12 tests conducted on 200 genotypes of poplar and oak, with 30% of SMPs containing 10 missing data points.

| | Method | Calculation Time (h:min:s) | RMSE | MAE | $R^2$ |
|---|---|---|---|---|---|
| Poplar | **Amelia II** | 00:04:50 | 0.078 | 0.053 | 0.959 |
| | **kNN** | 171:12:51 | 0.076 | 0.035 | 0.961 |
| | **LOCF** | **00:00:02** | 0.102 | 0.047 | 0.931 |
| | **LOESS** | 00:00:22 | 0.080 | 0.039 | 0.957 |
| | **methyLimp2** | 04:33:10 | 0.171 | 0.072 | 0.805 |
| | **MissForest** | 03:32:44 | **0.067** | 0.035 | **0.970** |
| | **NIPALS** | 05:10:05 | **0.067** | **0.034** | **0.970** |
| Oak | **Amelia II** | 00:01:07 | 0.084 | 0.059 | 0.958 |
| | **kNN** | 02:48:38 | 0.073 | **0.040** | 0.969 |
| | **LOCF** | **00:00:01** | 0.097 | 0.054 | 0.943 |
| | **LOESS** | 00:00:05 | 0.069 | 0.042 | 0.970 |
| | **methyLImp2** | 00:13:23 | 0.143 | 0.065 | 0.880 |
| | **MissForest** | 00:07:46 | 0.069 | 0.043 | 0.972 |
| | **NIPALS** | 00:43:55 | **0.067** | 0.042 | **0.973** |

Note: The results have been averaged over the 12 random replicas per percentage of missing data. Best results per metric are highlighted in bold for each dataset. RMSE: Root Mean Square Error. MAE: Mean Absolute Error. $R^2$ Coefficient of determination.

**Additional file: Table S2.** Overall CHH imputation performance was evaluated across 12 tests conducted on 200 genotypes of poplar and oak, with 30% of SMPs containing 10 missing data points.

| | Method | Calculation Time (h:min:s) | RMSE | MAE | $R^2$ |
|---|---|---|---|---|---|
| Poplar | **Amelia II** | 00:21:00 | 0.055 | 0.038 | 0.845 |
| | **kNN** | 25:07:58 | 0.043 | **0.023** | 0.898 |
| | **LOCF** | **00:00:16** | 0.060 | 0.032 | 0.899 |
| | **LOESS** | 00:01:40 | 0.044 | 0.025 | 0.901 |
| | **methyLImp2** | 145:04:03 | 0.061 | 0.031 | 0.807 |
| | **MissForest** | 28:02:42 | 0.044 | 0.025 | 0.899 |
| | **NIPALS** | 219:36:55 | **0.042** | 0.024 | **0.907** |
| Oak | **Amelia II** | 00:03:58 | 0.079 | 0.057 | 0.803 |
| | **kNN** | 19:52:06 | 0.069 | 0.042 | 0.861 |
| | **LOCF** | **00:00:03** | 0.088 | 0.053 | 0.751 |
| | **LOESS** | 00:00:25 | 0.063 | 0.040 | 0.871 |
| | **methyLImp2** | 05:07:22 | 0.084 | 0.047 | 0.773 |
| | **MissForest** | 01:58:22 | 0.063 | 0.041 | 0.876 |
| | **NIPALS** | 24:32:14 | **0.060** | **0.039** | **0.885** |

Note: The results have been averaged over the 12 random replicas per percentage of missing data. Best results per metric are highlighted in bold for each dataset. RMSE: Root Mean Square Error. MAE: Mean Absolute Error. $R^2$ Coefficient of determination. For kNN tests, a threshold was applied to retain only the first 15,000 SMPs before masking the data for imputation.
